# Variant intolerance scores in cattle

**DOI:** 10.64898/2026.07.28.741161

**Authors:** Simon Lanigan, Martijn FL Derks, Anna M Johansson, Martin Johnsson

## Abstract

Variant intolerance methods score the essentiality of genes based on large datasets of genetic variants and have been used in population genomics of humans and model organisms. In this paper, we estimated Residual Variation Intolerance Scores for protein-coding genes and predicted protein domains in cattle. In agreement with results from other species, the most variant-tolerant genes and domains included genes related to olfaction and adaptive immunity, whereas the least-variant tolerant genes and domains included genes involved in fundamental cellular processes. There was a moderate positive correlation with estimates from orthologous human genes. We provide estimates of variant intolerance for cattle may be useful for genomic analyses of deleterious variants and population genomics in cattle.

## Background

Genomic diversity carries traces of population genetic processes, including selection. Variant intolerance methods detect essential genes by their lack of functional variants in large sequence datasets [1–3]. They reflect selection against heterozygotes, that is, both the selection coefficients and dominance coefficients of new mutations [4]. Estimates of variant intolerance may be useful for interpreting genome-wide association studies, variant data such as putative loss-of-function alleles, or when targeting interventions such as genome editing and drugs towards genes and their products.

There are over 1.46 billion cattle worldwide (FAO, 2017), and an increasing amount are being genotyped as well as phenotyped. Cattle have been domesticated from two subspecies (*Bos taurus*) and (*Bos indicus*) which diverged 0.5 million years ago from extinct wild aurochs (*Bos primigenius*) [5]. DNA sequencing has provided a comprehensive genetic map of variation in the cattle genome which includes several million single nucleotide variants, thousands of insertion or deletion events alongside structural variants present in a genome of farm animals [6,7]. Investigations into the genomic diversity and functional genetic variation in cattle are relevant both for understanding cattle and their domestication, to help detect genetic disorders and defects and inform animal breeding and genetics, but also potentially for comparative studies with humans and other mammals. In order to support research in farm animal genetics and to enable comparative studies, it is useful to have estimates of population genomic statistics that are currently only available for humans and model organisms.

Variant intolerance scoring methods make use of genome-wide datasets of genetic variations to quantify selection on genes. Scoring genes based on the level of standing functional variation within a species was pioneered by Petrovski et al. [1] with the Residual Variant Intolerance Score (RVIS) method, which describes the effect of purifying selection on genes through a regression between the number of potentially functional variants observed in protein-coding sequence of a gene with the total number of variants, which serves as a proxy for mutational input. This method identifies genes that are relatively more or less tolerant to functional variation, suggesting their potential functional importance. Genes with a lower tolerance score are more likely to harbour functional mutation than their counterparts with higher scores. Other methods for variant intolerance scoring include probability of loss-of-function intolerance [3] and the missense tolerance ratio [8]. Similar logic can be applied to relevant parts of genes, such as exons or protein domains [2].

In this paper, we estimate residual variation intolerance scores of cattle genes, on the level of the gene and the predicted protein domain. We analyse variant-tolerant and intolerant genes using annotation term enrichment tests and compare the cattle scores to published scores from humans.

## Materials and methods

### Data

The variant dataset consisted of the publicly available run 9 of the 1000 Bull Genomes project [6], obtained in variant call file format from the European Nucleotide Archive (project accession PRJEB56689), including data from 1038 cattle. We filtered the variants based on the provided Variant Quality Score Recalibration [9,10] results, which uses multiple variant quality indicators in combination with known truth sets of variants to filter variant calls with the help of machine learning, by including only variants the “PASS” status. We used bcftools [11] to filter the variants, split multi-allelic sites, and to left-align insertion/deletion variants. Single nucleotide variants and insertion/deletion variants were considered together, because variant intolerance scoring aims at quantifying the total mutational input. 1000 Bull genomes variants were called on the ARS-UCD1.2 version of the cattle reference genome [12], and hence this version of the genome was used for all downstream analyses.

### Variant annotation

The Variant Effect Predictor (VEP) [13] was used to annotate variants as more or less likely to be functional using the VEP cache that includes Ensembl Genes version 107 as well as NCBI RefSeq gene annotation. We considered protein-coding variants classified by VEP as “HIGH” and “MODERATE” impact as potentially functional, e.g., stop and start gain and loss variants, frameshift variants, splice donor and acceptor variants, missense variants, in-frame indels. We also extracted protein-coding variants classified as unlikely to be functional (impact classification “LOW” according to VEP), e.g., splice region variants, start and stop retained variants and synonymous variants.

### Gene-based residual variation intolerance scoring

We used residual variation intolerance scores (RVIS) method [1], which scores protein-coding genes based on their amount of common potentially functional variation compared to the overall number of observed variants in the gene. For the total number of functional variants, we considered variants classified by VEP as “HIGH” and “MODERATE”, and the total variant count included those variants plus variants classified by VEP as “LOW” impact.

We used vcftools [14] (version 0.1.16) to estimate allele frequency. Allele frequencies and variant annotation were combined to count the number of potentially functional variants and the total number of protein-coding variants in each gene. The analysis was performed at the level of the gene, and thus we included all variants that were annotated as potentially affecting any of the transcripts associated with the gene and counting each variant only once if it overlapped multiple transcripts of the same gene.

The threshold for considering a potentially functional variant “common” was set at an alternate allele frequency of 0.1%, following the original authors of the RVIS method [1]. Variants below this threshold were considered rare and thus contributed to the total number of variants in the gene, but not to the number of common functional variants. To evaluate the impact of the choice of allele frequency threshold, we also performed counts with no allele frequency filtering and with a threshold of 1% alternate allele frequency.

A simple linear regression was fitted with the number of common functional variants per gene as response variable and the total number of variants in each gene as predictor. The residual variation intolerance score for each gene is the studentized residual from this linear model. The linear model was performed in the R statistical environment [15] with the base R lm function, and the studentized residuals were calculated with the studres function of the MASS package.

We performed the analysis separately for the Ensembl and NCBI gene annotation. For the main analysis, we present results from the Ensembl annotation. We included only genes that had at least one protein-coding variant detected.

### Clustering of genes with high and low scores

To detect clustering of genes with high and low RVIS, we divided the genome into windows of 1 Mbp using the GenomicRanges R package [16] and counted the number of genes in each window. The genomic locations of genes were taken from Ensembl Gene annotation.

In order to generate an empirical null distribution reflecting a lack of clustering, we sampled an equal number of randomly selected genes from Ensembl Genes that had estimated RVIS values. We repeated this random sampling 1000 times, counted the number of genes overlapping 1-Mbp windows and recorded the maximum number from each replicate. We took the 95% quantile of this distribution as the threshold for significant genomic clustering.

### Domain-based variant intolerance scoring

We also estimated domain-based residual variation intolerance scores [2] for protein domains. The domain annotation used was predicted Pfam domains [17] from the Ensembl Genes database, accessed through BioMart [18]. For each gene, we scored domains only on the principal transcript as assigned in APPRIS transcript annotation [19], in order to avoid pseudoreplication arising from counting the same domain multiple times.

The scores were estimated analogously to gene-based RVIS, by counting the total number of variants and the number of common potentially functional variants falling within each domain. As before, variants classified as “HIGH” or “MODERATE” impact by VEP were considered potentially functional, and the cutoff for a variant to be considered common was 0.1%. To evaluate the impact of the choice of allele frequency threshold, we also performed counts with no allele frequency filtering and with a threshold of 1% alternate allele frequency.

As before, we included only variants that had at least one protein-coding variant detected. We fit a simple linear regression with the number of common functional variants in each domain as response variable and the total number of variants in the domain as predictor and extracted domain RVIS scores as studentized residuals.

### Enrichment testing

To detect enriched annotation terms among the most and least variant-tolerant genes, we extracted genes with the top and bottom 10% of RVIS scores from Ensembl genes. The top and bottom genes were then analysed further using the Gene Ontology and GALLO for quantitative trait locus overlaps, and the top and bottom domains were analysed for Pfam domain enrichment. For enrichment analysis of Gene Ontology terms [20,21] and Pfam domains, we performed term enrichment tests using Fisher’s Exact Test in R, applying Holm correction to the p-values to account for multiple testing. The reference list for this analysis consisted of all cattle genes or domains for which we estimated RVIS. For enrichment analysis of quantitative trait loci, we used the R-package GALLO [22] and the QTL from the Cattle QTLdb database version 58 [23]. We extracted the position of the genes with top and bottom 10% scores on the ARS-UCD1.2 genome from Ensembl. We applied Holm correction to the p-values to account for multiple testing.

### Comparisons to human orthologs

For comparison to human orthologs, scores were aligned with publicly available human RVIS estimates [24] for orthologous human genes. As recommended by the authors, we used the human RVIS estimated from people of African ancestry. We used BioMart to access Ensembl comparative data [25] to identify one-to-one orthologous genes and estimated the Spearman rank correlation of RVIS between species. To identify the genes that differ the most in their intolerance score between species, we ranked the genes based on their estimated human and cattle RVIS, and found genes with the largest positive and negative rank differences.

## Results

### Gene-based intolerance scores

Residual variation intolerance scores (RVIS) [1] were estimated to assess the intolerance to variation among cattle genes. The method uses a simple linear model with the number of common potentially functional variants in a gene as response variable and the total number of variants in the gene as predictor, to estimate constraint on protein-coding genes. The intolerance scores are studentized residuals of this model, where a low score represents a gene that is relatively intolerant to variation. In total, 21,841 genes were scored based on the Ensembl Genes database and potentially protein-coding variants from the public dataset from the 1000 Bull Genomes project run 9. Figure 1 shows the regression between number of common functional variants and the total number of variants, highlighting the top and bottom 10% genes. The estimated RVIS values ranged from -9.81 to 12.9. Table 1 shows the top and bottom 10 genes. The top 10 included *MKI67* (*marker of proliferation Ki- 67*) as the most variant-tolerant, and 5 novel genes. The bottom 10 included *TRRAP* (*transformation/transcription domain associated protein*) as the most variant-intolerant, as well as ryanodine receptors and dynein genes.

**Figure 1.**
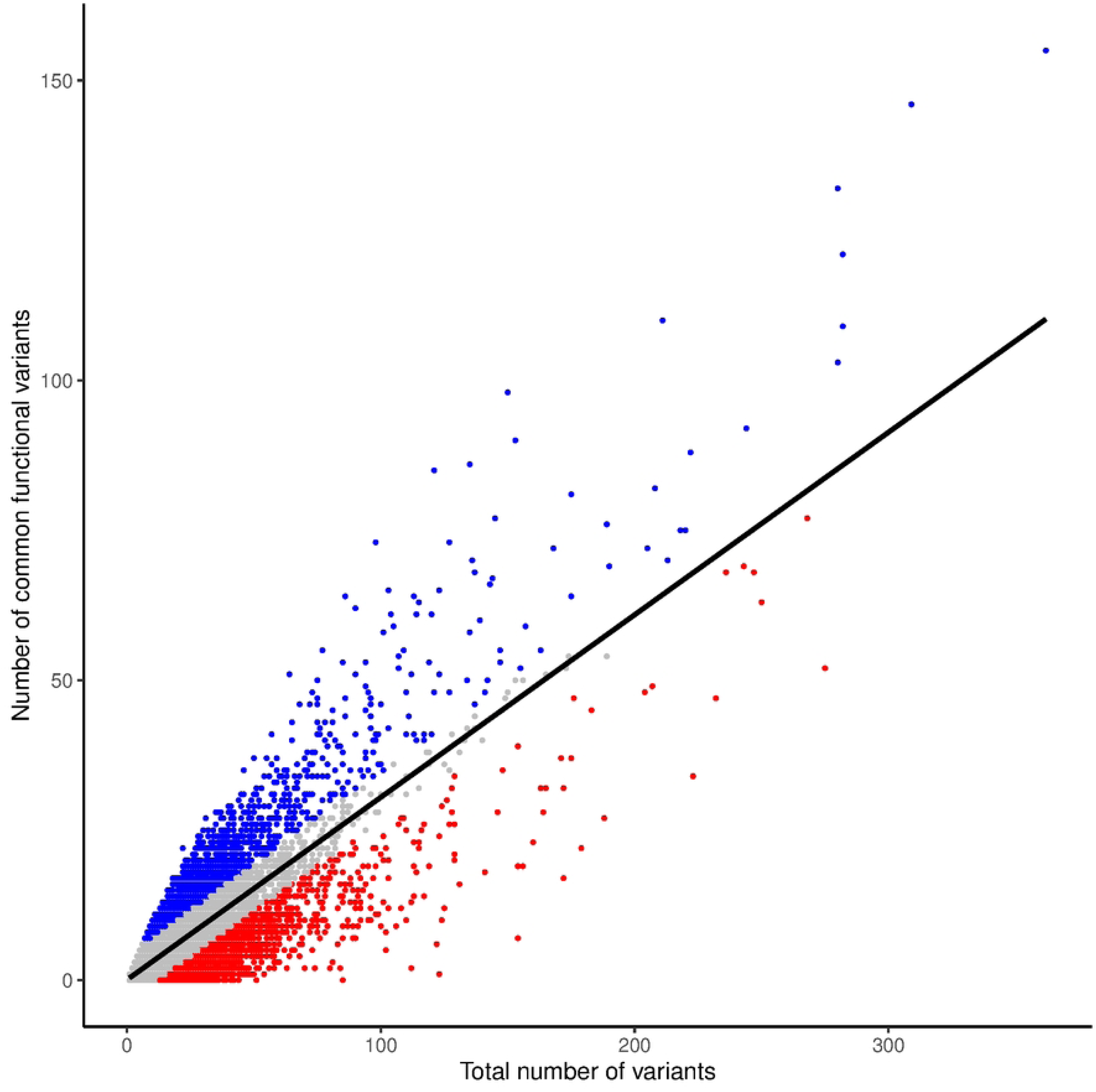
Gene-based RVIS. Scatterplot of the total number of variants and the number of common potentially functional variants in Ensembl Genes. The regression line represents the simple linear regression used to estimate RVIS. Red points correspond to genes with bottom 10% RVIS values and blue points correspond to genes with top 10% RVIS values.

**Table 1.**
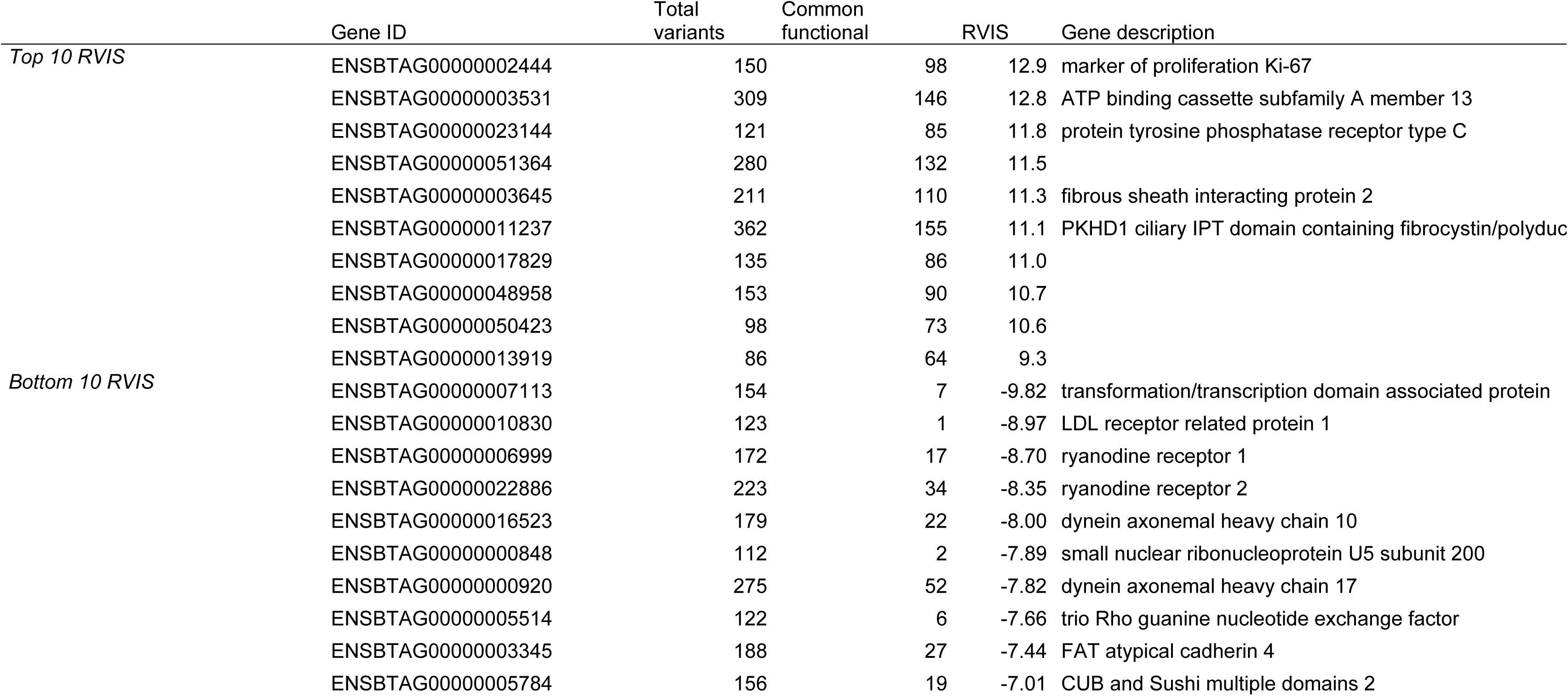
Top and bottom ten genes with the highest and lowest estimated RVIS, with Ensembl Gene identifier, total number of variants, number of common potentially functional variant, RVIS and gene description from the Ensembl Gene database.

The residual variation intolerance method has one parameter that may be adjusted, namely the threshold for considering a variant common or rare. In addition to the main analysis using the threshold of 0.1% allele frequency used by the original authors [1], we also estimated intolerance scores using no frequency filtering or a threshold of 1% (S2 Figure). The Spearman correlation of was 0.94 between scores estimated with the standard threshold and no filtering, and 0.92 between scores with standard threshold and 1%.

In addition to the main analysis using the Ensembl gene annotation, we also performed the analysis based on the NCBI gene annotation, the results of which are shown in S1 Figure and S1 Table. The complete variant intolerance scores for all genes are available as S1 Data (Ensembl genes) and S2 Data (NCBI genes).

The most variant-intolerant genes were relatively evenly distributed in the genome, whereas for the most variant-tolerant genes there were some genomic clusters.

Figure 2 shows the genomic distribution of genes with top and bottom 10% RVIS values and counts of top and bottom genes in 1-Mbp windows of the genome. We tested by clustering by randomly sampling genes and counting the number of genes in 1-Mbp windows, giving an empirical null distribution. We identified 10 windows that passed the 95% threshold. The cluster containing the most top 10% variant-tolerant genes was located on chromosome 23 at 27-31 Mbp, overlapping the MHC and an olfactory receptor gene cluster.

**Figure 2.**
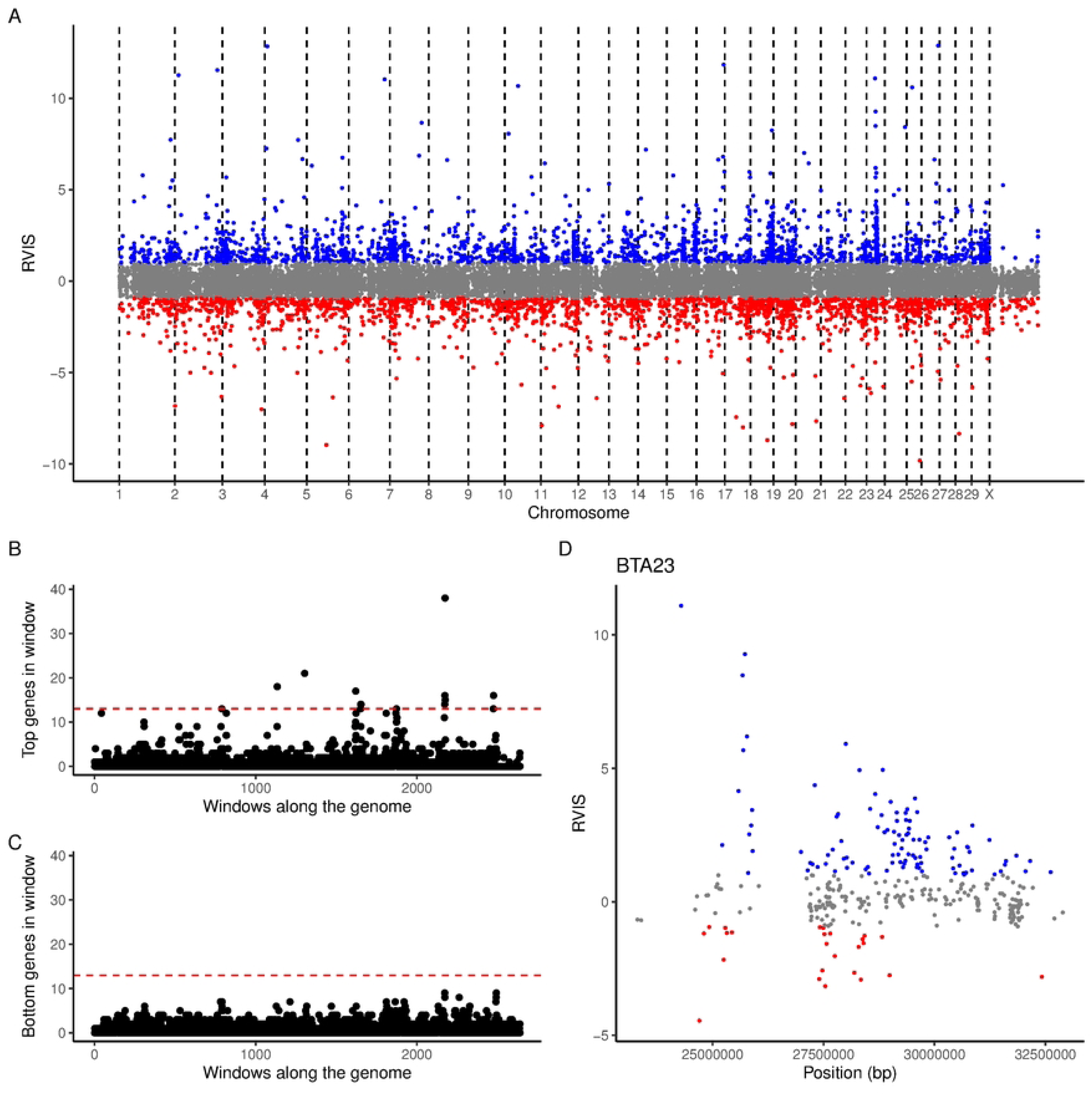
Genomic position of scored RVIS genes. Panel A shows genomic positions of all scored genes with their RVIS values. Red points correspond to genes with bottom 10% RVIS values and blue points correspond to genes with top 10% RVIS values. Panels B and C show the count of top and bottom genes in 1-Mbp windows along the genome. The horizontal dashed line shows a significance threshold for clustering, corresponding to the 95% percentile of simulated clustering. Panel D shows a zoomed-in view of the region on chromosome 23 around the cluster with the highest number of top 10% genes.

**Figure 3.**
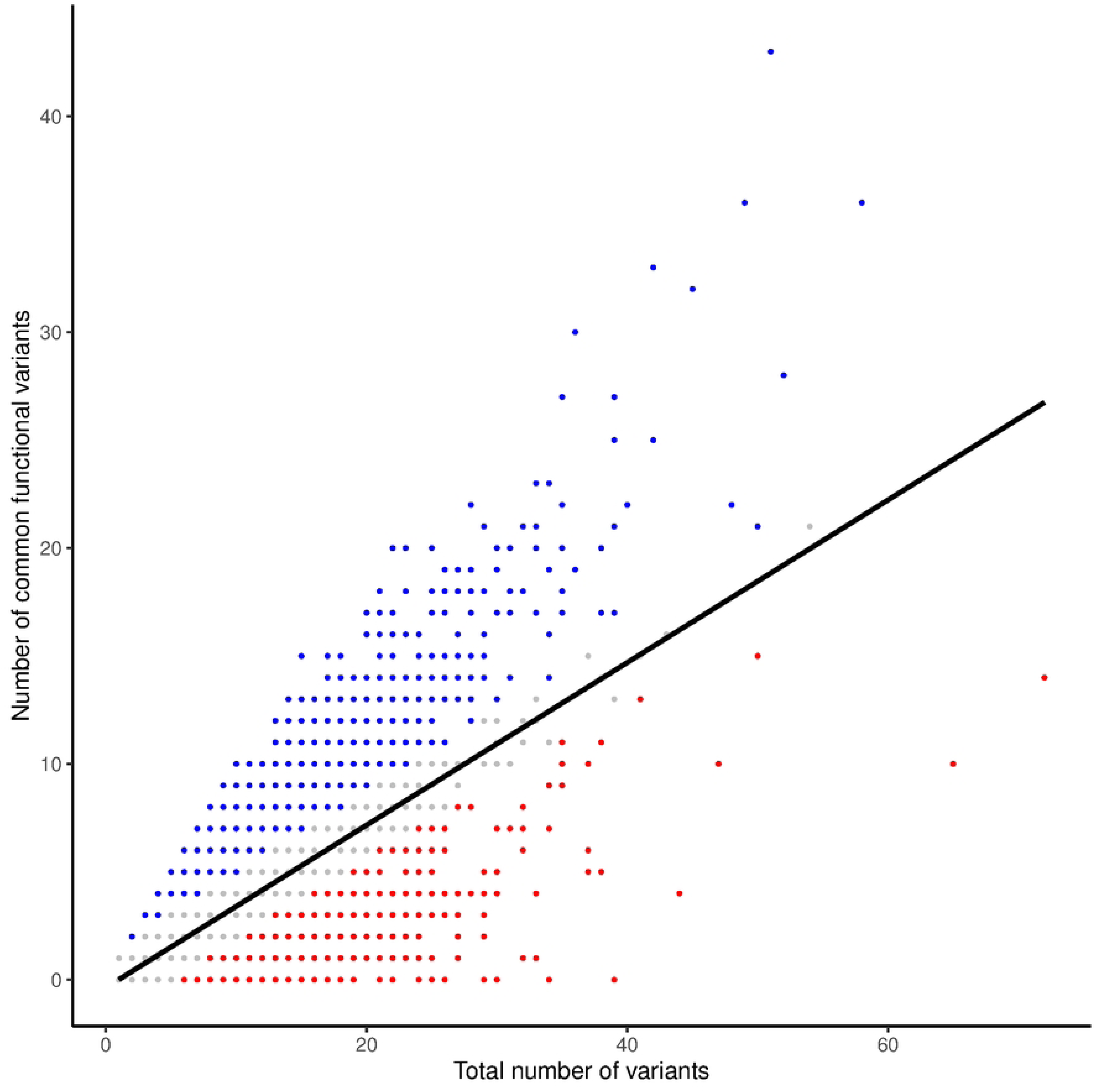
Domain-based RVIS. Scatterplot of the total number of variants and the number of common potentially functional variants in Pfam domains. The regression line represents the simple linear regression used to estimate domain RVIS. Red points correspond to domains with bottom 10% domain RVIS values and blue points correspond to domains with top 10% domain RVIS values.

To interpret the scores, we performed annotation term enrichment of the most and least variant-tolerant genes. Enriched annotation terms among the most variant-tolerant genes include terms related to olfaction, whereas enriched annotation terms among the least variant-tolerant genes include overrepresentation of fundamental cellular processes (S2 Table). Further, we overlapped the top and bottom 10% genes with QTL from the Cattle QTLdb database and tested for QTL enrichment with the GALLO package. The significantly enriched traits are shown in S3 Table.

### Domain-based intolerance scores

Following Gussow et al. [2], we also estimated variant intolerance scores at the level of protein domains, using predicted protein domains from the Pfam database. The method is analogous to gene-based intolerance scoring, applied to functional meaningful partitions of genes. Figure 2 shows the regression between number of common functional variants and the total number of variants, highlighting the top and bottom 10% domains. Table 2 shows the top and bottom 10 domains, including class II MHC antigen domains and olfactory receptors among the most variant-tolerant domains, and among others myosin and heat shock protein domains as the most variant-intolerant. S4 Table shows significantly enriched domain families among the top and bottom 10% domains. The complete domain RVIS estimates for all domains are available in S3 Data.

**Table 2.**
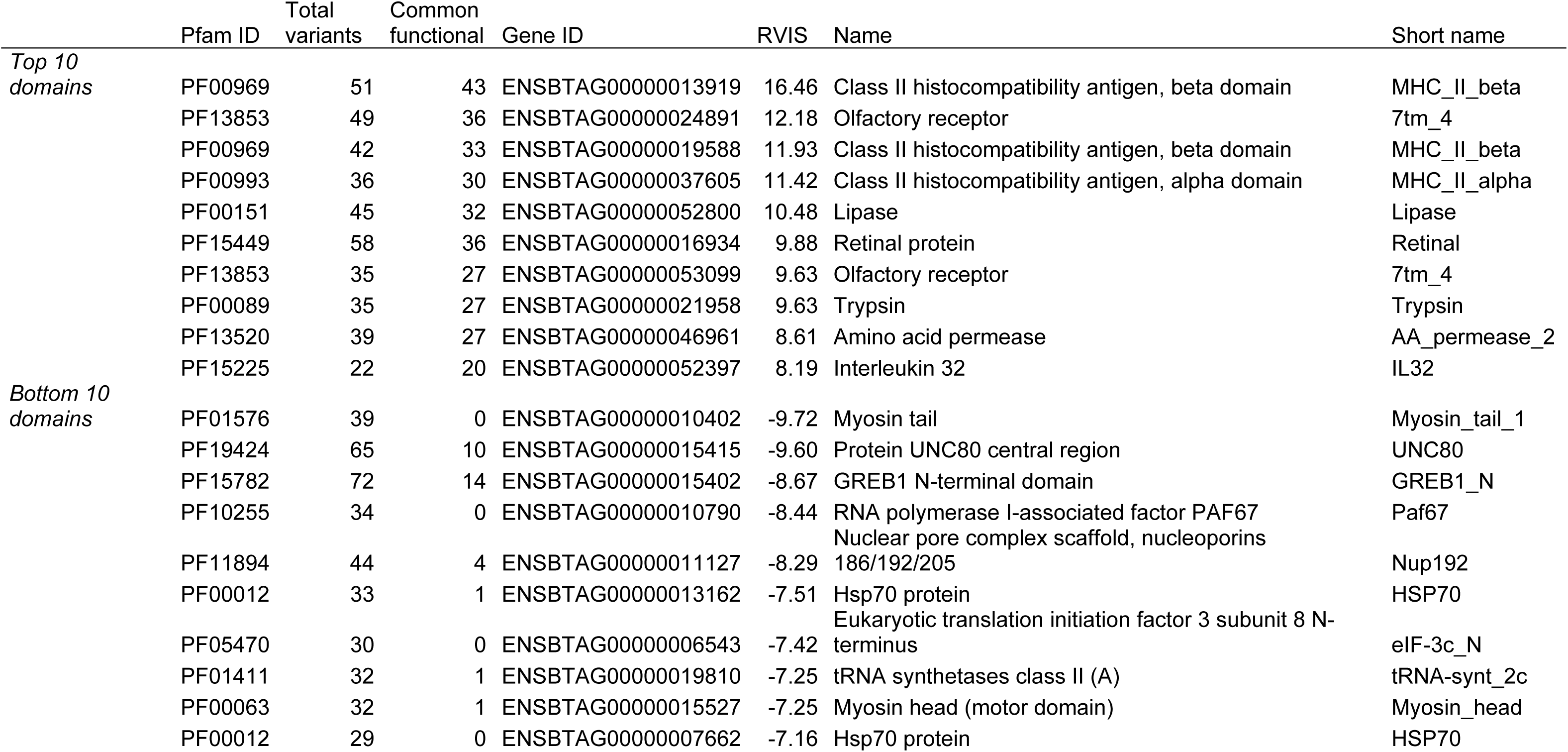
Top and bottom ten domains with the highest and lowest estimated domain RVIS, with Pfam identifier, total number of variants, number of common potentially functional variant, Ensembl Gene identifier, RVIS and domain names from the Interpro database.

As with the gene-based analysis, we also performed additional analyses using no frequency filtering or a threshold of 1% (S2 Figure). The Spearman correlation of was 0.91 between scores estimated with the standard threshold and no filtering, and 0.89 between scores with standard threshold and 1%.

### Comparison to human orthologs

We compared our gene-based variant intolerance scores from cattle to estimates from humans published by Han et al. [24], using one-to-one orthologous genes. Figure 4 shows a scatter plot of RVIS values for human versus bovine genes. The Spearman rank correlation coefficient between the human and cattle RVIS was 0.49.

**Figure 4.**
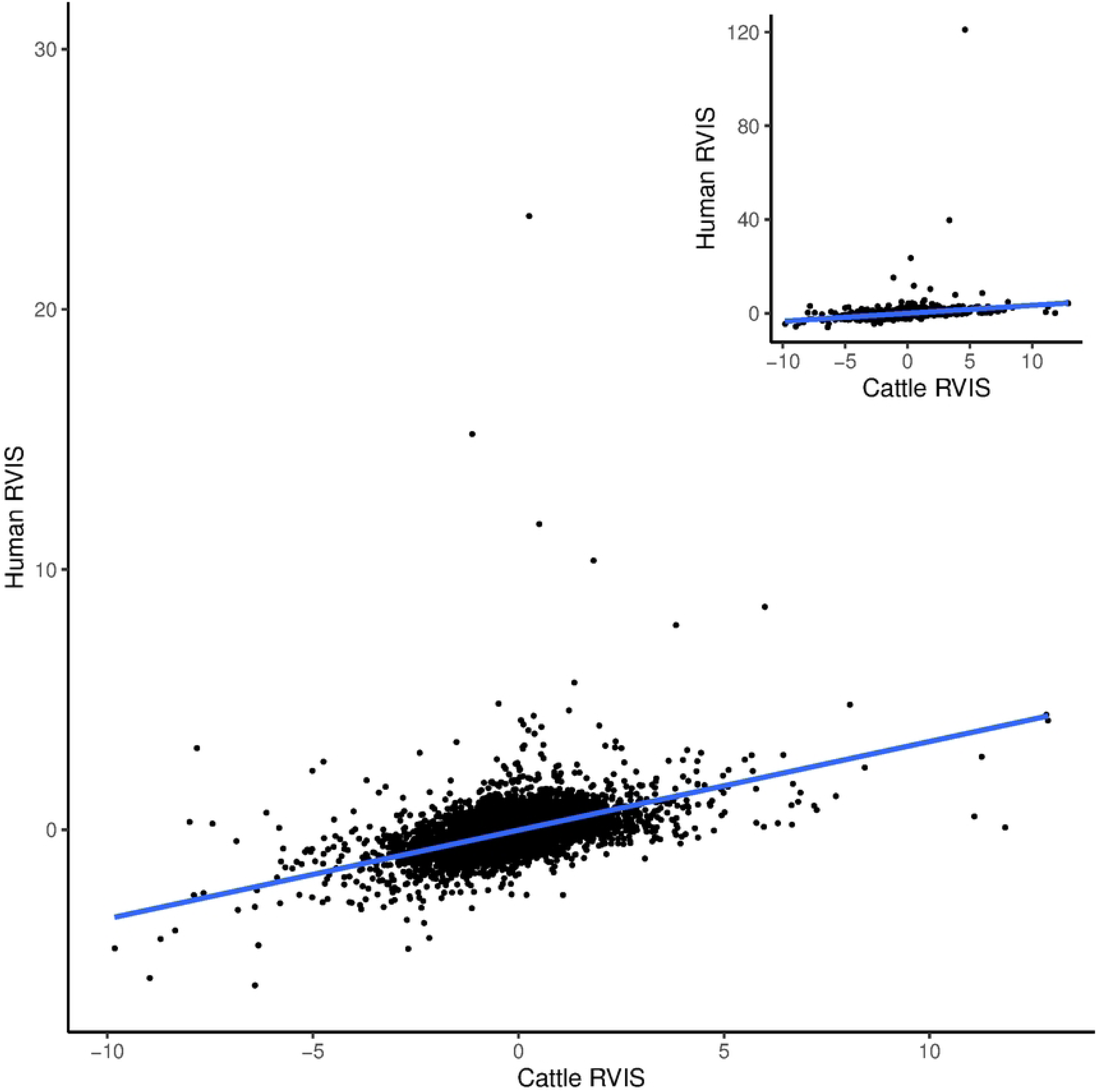
Comparison to human orthologues. Scatterplot of cattle RVIS compared to human RVIS [24] on 1-to-1 orthologues. The main plot has been zoomed in to exclude two human genes with extremely high RVIS values, with the full scatterplot shown in the top right corner. The blue line represents a simple linear regression.

We identified genes that differ the most in their intolerance between species, in the sense that they show the greatest difference in rank between human and bovine RVIS. The most reranked genes that are more variant-tolerant in cattle than include several genes related to neuronal function, whereas genes that are less variant- tolerant in cattle include muscle-related genes (Table 3). The top 10% most reranked genes that were more variant-tolerant in cattle were enriched for terms related to intracellular signalling and voltage-gated calcium channels whereas the top 10% most reranked genes that were more variant-intolerant were enriched for terms related to the cytoskeleton and microtubuli (S5 Table).

**Table 3.** The top 10 most reranked genes that were more or less variant-intolerant, respectively, in cattle compared to humans, with cattle Ensembl Gene identifier, human Ensembl Gene identifier, gene symbol, cattle RVIS, and human RVIS.

|  | Cattle Gene ID | Human Gene ID | Symbol | Cattle RVIS | Human RVIS |
| --- | --- | --- | --- | --- | --- |
| *More tolerant in cattle* | ENSBTAG00000020238 | ENSG00000079841 | *RIMS1* | 3.08 | -1.11 |
|  | ENSBTAG00000045610 | ENSG00000100346 | *CACNA1I* | 1.31 | -1.36 |
|  | ENSBTAG00000017618 | ENSG00000130758 | *MAP3K10* | 2.14 | -0.94 |
|  | ENSBTAG00000020252 | ENSG00000169992 | *NLGN2* | 1.59 | -1.09 |
|  | ENSBTAG00000011581 | ENSG00000164061 | *BSN* | 1.08 | -2.51 |
|  | ENSBTAG00000009714 | ENSG00000004777 | *ARHGAP33* | 1.29 | -1.12 |
|  | ENSBTAG00000015170 | ENSG00000115850 | *LCT* | 1.37 | -0.97 |
|  | ENSBTAG00000017349 | ENSG00000240184 | *PCDHGC3* | 1.39 | -0.95 |
|  | ENSBTAG00000024199 | ENSG00000055609 | *KMT2C* | 1.17 | -1.05 |
|  | ENSBTAG00000011102 | ENSG00000186815 | *TPCN1* | 1.18 | -1.01 |
| *More intolerant in cattle* | ENSBTAG00000000920 | ENSG00000187775 | *DNAH17* | -7.82 | 3.13 |
|  | ENSBTAG00000016784 | ENSG00000118997 | *DNAH7* | -4.74 | 2.62 |
|  | ENSBTAG00000006907 | ENSG00000183091 | *NEB* | -5.01 | 2.26 |
|  | ENSBTAG00000009387 | ENSG00000036448 | *MYOM2* | -3.69 | 1.90 |
|  | ENSBTAG00000020657 | ENSG00000083857 | *FAT1* | -3.23 | 1.65 |
|  | ENSBTAG00000009331 | ENSG00000160111 | *CPAMD8* | -3.40 | 1.43 |
|  | ENSBTAG00000017434 | ENSG00000185038 | *MROH2A* | -2.40 | 2.96 |
|  | ENSBTAG00000018560 | ENSG00000158486 | *DNAH3* | -2.81 | 1.23 |
|  | ENSBTAG00000020243 | ENSG00000165124 | *SVEP1* | -2.16 | 1.44 |
|  | ENSBTAG00000015526 | ENSG00000184571 | *PIWIL3* | -1.65 | 1.89 |

## Discussion

In this paper, we estimated residual variation intolerance scores (RVIS) for genes and protein domains in cattle based on genetic variants from the 1000 Bull Genomes project. A total of 21,841 genes were scored, and further investigation focused on the top and bottom genes to detect biological themes among the most and least variant- tolerant genes in the cattle genome.

A low RVIS value suggests that a gene is intolerant to functional variation, likely due to purifying selection, as intolerance scores reflect selection against new variants in a heterozygous state [4]. A high RVIS value suggests that a gene is tolerant to functional variation, which may be due to relaxed purifying selection or balancing selection that favours diversity. The scores are based on standardised residuals from a genome-wide regression between the number of common potentially functional variants and the total number of variants, serving as a proxy for mutational input.

Genes with lower RVIS scores are often associated with essential biological functions and are more conserved [1,2].

In our analysis, the most variant-intolerant genes and domains included fundamental cellular functions such as protein phosphorylation, signal transduction and cell adhesion, as well as rhodopsin and the cytoskeleton. The most variant-tolerant genes and domains included functions related to olfaction and immunity. Variant- intolerant genes were relatively uniformly spread across the genome, whereas there was some genomic clustering of the most variant-tolerant genes. Overlap and enrichment of published quantitative trait loci suggested that both variant-tolerant and variant-intolerant genes were enriched for regions associated with quantitative traits in cattle. Comparing intolerance scores between cattle and human orthologs revealed a moderate positive correlation. We will now discuss some of the most and least intolerant genes, illustrating their potential function, potential implications for cattle genetics, and finally limitations and assumptions that have not been discussed previously.

### The function of the most and least variant-tolerant genes

The gene with the highest RVIS in our dataset was *MKI67* (*ENSBTAG00000002444*), which encodes a protein involved in chromatin organisation [26] that is used as a cell proliferation marker particularly for cancer. Its human homolog is also scored as highly variant-tolerant [24]. Other highly variant-tolerant genes include ABC-transporter gene *ABCA13* (*ENSBTAG00000003531*)*, PTPRC* (*ENSBTAG00000023144*) which encodes a protein tyrosine phosphatase involved in immune function, *FSIP2* (*ENSBTAG00000003645*) which is involved in sperm function, and *PKHD1* (*ENSBTAG00000011237*) involved in function of primary cilia. Some of genes with the highest RVIS were novel genes without associated symbols. Their extremely high scores may suggest that they are poorly annotated and not in fact functional protein-coding genes. One of them, however, *ENSBTAG00000013919* is located among known major histocompatibility complex (MHC) genes on chromosome 23 and may be an immune gene, based on it being mapped as an orthologue of mouse *histocompatibility 2, class II antigen E beta*.

Immune genes involved in antigen presentation such as the genes of the MHC are expected to show high variant intolerance scores (i.e., to be scored as variant-tolerant) due to balancing selection, and appear among genes with high RVIS in humans [1,24]. At the same time, the MHC region is structurally complex, repetitive, and structurally variable [27]. Therefore, both genome assembly and variant calling from short read sequencing are difficult, and it is unlikely that the gene annotation and variant calling of MHC genes is complete and accurate. In our data, we detect highly variant-tolerant genes and domains related to the adaptive immune system, as well as an enrichment of MHC class I-related and immunoglobulin protein domains in the domain RVIS analysis and several MHC class II domains scored as the top 10 most variant-tolerant. However, in the gene-level RVIS analysis, we do not detect an enrichment of MHC-related GO terms. On the genome-level, we find clustering of highly variant-tolerant genes on chromosome 23 in the region of the cattle MHC.

Olfactory receptor genes were also observed among the most variant-tolerant genes, a phenomenon reported already in the first genome-wide investigations of SNP heterozygosity and variant intolerance in humans [1,28]. In our data, we found enrichments of multiple Gene Ontology terms related to olfaction among the most variant-tolerant genes, and the most strongly enriched protein domain family was the 7-transmembrane domain associated with olfactory receptor genes. The cluster of highly variant-intolerant genes on chromosome 23 also overlap an olfactory receptor gene cluster which is located adjacent to the MHC. Olfactory receptor genes are subject to rapid gene family evolution, including expansion and contraction in mammals [29,30]. We do not know whether the high variant-tolerance is due to balancing selection like the MHC or due to this rapid diversification. Variant calling from short reads may also be limited due to the repetitive nature of these gene clusters. Pan-genome efforts and long read sequencing may be necessary to accurately estimate variant intolerance in regions such as the MHC and olfactory receptor gene clusters.

Conversely, essential genes and domains under strong selective constraint are expected to have low variant intolerance scores, reflecting less segregating functional variation than expected from the mutational input. The gene with the lowest RVIS in our data is *TRRAP* (*ENSBTAG00000007113*), encoding part of a histone acetyl transferase complex [31]. Its human homolog is also scored as highly variant-intolerant [24]. Notably, ryanodine receptors *RYR1* (*ENSBTAG00000006999*) and *RYR2* (*ENSBTAG00000022886*) are both among the ten most variant-intolerant genes. Along with enrichment of Gene Ontology terms related to calcium-ion transport and calcium ion binding, they suggest that genes involved in calcium regulation are highly constrained. Enriched Gene Ontology terms also involve broad terms associated with other fundamental biological processes and structures such as protein phosphorylation, signal transduction, the cytoskeleton and even a significant enrichment of protein binding, which is hard to interpret because of its commonality, associated with thousands of genes. In contrast to the olfactory genes with high RVIS, the most enriched domain among those with low RVIS was the 7- transmembrane domain associated with rhodopsin, suggesting that this function involved in sight is highly constrained. Genetic variation in rhodopsin affects rhodopsin stability and retinal disease [32].

### Potential implications for cattle genetics

Variant intolerance scores have found use in evolutionary analyses of development [33], in variant prioritisation for genetic disorders [1–3,34,35], and in analyses of the genetic architecture of complex traits [36,37]. This suggests that variant intolerance scoring in farm animals may be useful in screening for deleterious variants and genetic defects, where a potential loss of function variant in a variant-intolerant gene might be more likely to be deleterious than a potential loss-of-function variant in a gene that is scored as variant-tolerant. Analyses of genetic architecture in humans [37] suggests that variant-intolerant genes are more likely to also contain variants with larger effect sizes for complex diseases. Therefore, one may speculate that variant intolerance scoring could be part of weighting variants for genomic prediction.

In our enrichment analysis of published quantitative trait loci relating to cattle traits, both sets of highly variant-tolerant and variant-intolerant genes were enriched in regions associated with production and reproduction traits in cattle. The list of enriched traits showed more meat and carcass traits (e.g., bone weight, carcass weight and muscle area) for high RVIS genes and more milk-related traits (e.g., milk fat yield, milk fat percentage) for low RVIS genes. However, given the very large number of overlaps, and the fact that the associated regions often cover many genes, the genomic resolution of this analysis is poor, making it hard to interpret. To investigate the association between variant intolerance and complex traits in cattle, one might need more granular approaches, such as fitting genome partitioning models to quantify the genetic variance associated with genes with low or high scores, analogous to heritability partitioning studies using functional annotation [38,39].

Investigating shared genes helps identify evolutionarily conserved genes. The moderate positive correlation in RVIS scores might indicate some similarities in the evolutionary constraints or functional importance of genes between humans and cattle. However, there are clearly differences between the species. For example, our results suggest that dynein genes are under stronger constraint in cattle than in humans, whereas voltage-gated calcium channel activity, including several genes active in the central nervous system, are under stronger constraint in humans.

Comparative studies of variant intolerance between closely related species such as domestic and wild ruminants may illuminate the evolution of variant intolerance.

Further, it is an open question how much intolerance scores are affected by recent changes in selection pressures such as during domestication or local adaptation. Future research may address population differences, using datasets with more even representation of subspecies and breeds of cattle.

### Limitations and assumptions

The study is limited by the resolution of variant calling from short-read sequencing, gene annotation, and the sampling of worldwide cattle diversity.

When identifying highly variant-tolerant genes, there is a risk of scoring genes high because of annotation or variant calling errors. For example, a pseudogene erroneously annotated as a functional protein-coding gene would likely score high, because it may freely accumulate segregating variant within its open reading frames, that may appear as potentially functional in variant effect prediction. As mentioned, the list of top 10 most variant-tolerant genes included several novel genes from the Ensembl Gene database, i.e. gene models that have less support from previous known genes. It is possible that some of them are due to errors in gene annotation, and some caution is warranted when interpreting genes and domains with high RVIS values for this reason.

Genetic variant datasets based on short-read sequencing, such as the 1000 Bull genomes, are limited to single nucleotide variants and short insertion/deletions. Longer structural variants are entirely missing, and the power to detect short insertion/deletions is limited [40]. Further, highly repetitive and structurally variable regions, including the olfactory receptor gene clusters and the major histocompatibility complex, are particularly challenging for short-read sequencing and variant calling. Again, caution is warranted when interpreting results from these regions, as there are likely many variants that are missed. With the increasing availability of long-read sequence data from cattle, more accurate estimates will be possible [41].

This analysis used the whole public part of the 1000 Bull genomes dataset, run 9, jointly, as large publicly available sample of worldwide cattle diversity. However, with larger samples of breeds and of the *taurus* and *indicus* subspecies, it would be possible to estimate stratified variant intolerance scores, and investigate to what extent recent changes in selection pressure due to domestication and breeding affect variant intolerance.

## Supporting information captions

S1 Figure. Gene-based RVIS calculated on NCBI genes. Scatterplot of the total number of variants and the number of common potentially functional variants in NCBI annotation. The regression line represents the simple linear regression used to estimate RVIS. Red points correspond to genes with bottom 10% RVIS values and blue points correspond to genes with top 10% RVIS values.

S2 Figure. Sensitivity of RVIS values to the threshold for separating common from rare variants. The panels show scatterplots with Spearman rank correlations for RVIS estimated with no allele frequency filtering, the standard 0.1% frequency threshold or a 1% frequency threshold, as well as histograms of the distributions of scores. The top panels show gene-based scores and the lower panels domain-based scores.

S1 Table. Top and bottom ten genes with the highest and lowest estimated RVIS in the NCBI annotation, with gene identifier, total number of variants, number of common potentially functional variant and RVIS.

S2 Table. Significantly enriched Gene Ontology terms, showing the GO term accession, number of genes annotated with that term in the top or bottom 10%, number of genes annotated with that term among all scored genes, Holm corrected p-value, and the GO term name.

S3 Table. QTL enrichment results showing QTL trait, number of QTL overlapping the top or bottom 10% genes, Holm-corrected p-value, and trait type classification.

S4 Table. Domain enrichment. Significantly enriched Pfam domain families, showing Pfam identifier, number of domains of that family in the top or bottom 10%, number of domains of that family with that term among all scored domains, Holm corrected p- value, and the Pfam domain names.

S5 Table. Significantly enriched Gene Ontology terms among the 10% most reranked genes that were more or less variant-intolerant, respectively, in cattle compared to humans. The columns show the GO term accession, number of genes annotated with that term in the top or bottom 10%, number of genes annotated with that term among all scored genes, Holm corrected p-value, and the GO term name.

S1 Data. All RVIS estimates from Ensembl genes.

S2 Data. All RVIS estimates from NCBI genes.

S3 Data. All domain RVIS estimates.

## Notes

### Competing Interest Statement

The authors have declared no competing interest.

